# Protocol for Seahorse 3D Mito Stress assay in patient-derived pediatric brain tumor single neurospheres

**DOI:** 10.64898/2025.12.03.691986

**Authors:** Stefania Tocci, Jessica W. Tsai

**Affiliations:** Children’s Hospital Los Angeles, Cancer and Blood Disease Institute, Los Angeles, California, United States; Children’s Hospital Los Angeles, Saban Research Institute, Los Angeles, California, United States; Keck School of Medicine of University of Southern California, Department of Pediatrics, Los Angeles, California, United States; USC Norris Comprehensive Cancer Center, Epigenetic Regulation in Cancer Program, Los Angeles, California, United States

## Abstract

Metabolism is essential for cellular functions and is often altered in cancer. While Seahorse assays are well established for measuring metabolic changes in 2D cell cultures, their application to 3D models remains challenging. Here, we present a step-by-step protocol for plating individual brain tumor neurospheres in the assay microplate to measure their bioenergetics. We provide actionable recommendations and highlight pitfalls to avoid. This approach enables exploration of dynamic metabolic changes in patient-derived brain tumor neurospheres following genetic or pharmacologic interventions.

**Graphical abstract:** 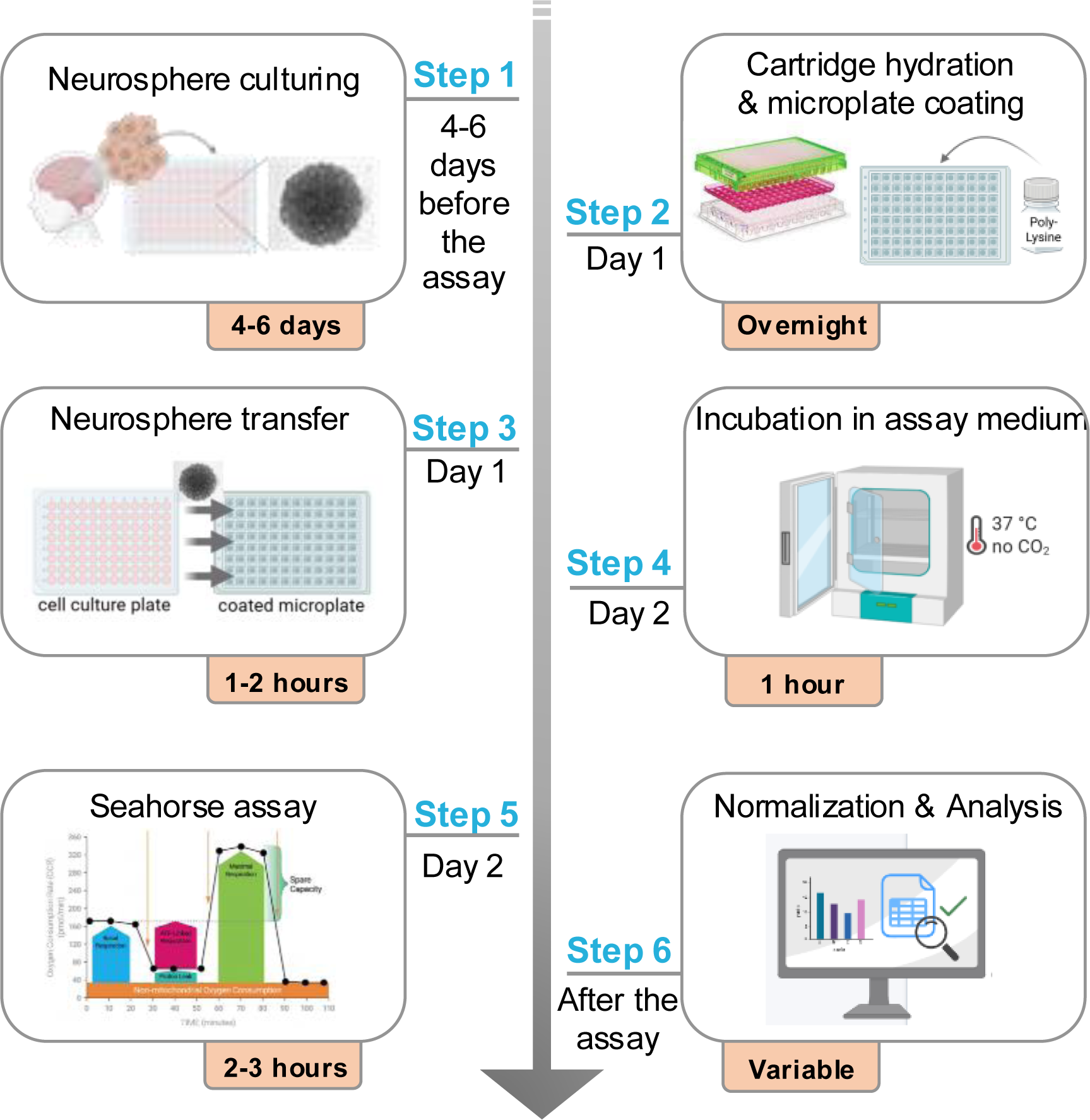

**Highlights:**

- Optimization of Seahorse assay protocol for single patient-derived brain tumor spheroids
- Step-by-step instructions for successfully re-plating and transferring single brain tumor spheroids
- Importance of microplate coating for the successful execution of Seahorse assays
- Imaging of the microplate before and after the assay allows for normalization of metabolic parameters based on neurosphere size

## Before you begin

Cancer cells rewire their metabolism to sustain rapid growth, adapt to environmental stress, and evade regulatory mechanisms^1–3^. This metabolic reprogramming creates vulnerabilities that can be leveraged therapeutically. Pediatric brain tumors also display distinct metabolic characteristics, yet comprehensive profiling of these alterations is limited^4–6^. Characterization of metabolic dynamics is crucial for understanding pediatric tumor biology and identifying potential therapeutic targets. The neurosphere model used in this protocol consists of 3D floating, spherical clusters of cells derived from pediatric brain tumors. The spheroids originate from cancer stem cells, thus maintaining the phenotype and molecular characteristics of the original tumor, providing a valuable system for studying metabolic alterations^7–9^. Neurospheres are typically cultured in serum-free conditions, thus preserving their self-renewing properties. In contrast, serum-containing media used in 2D monolayer cultures induces and promotes cell differentiation and significant metabolic changes^10^. While the neurosphere model is an *in vitro* cell model with inherent limitations, it offers a more tractable and reproducible alternative to *in vivo* models, such as animal systems, where assessing metabolic dynamics can be technically challenging and cost prohibitive.

In this protocol, we describe how to study metabolic responses in single pediatric brain tumor neurospheres. While our approach focuses on the Mito Stress assay, it can be adapted to studying other aspects of cellular metabolism, including glycolysis, fatty acid synthesis, and oxidative stress. By using a 3D patient-derived model, this optimized protocol provides a platform to measure metabolic changes that more accurately reflect tumor physiology.

## Innovation

Seahorse assays are extensively employed to investigate cellular metabolism in 2D monolayer cultures. While existing protocols have been adapted for 3D models such as organoids^11^, they often rely on modifications to 2D methodologies and lack specificity. Additionally, studies on metabolic profiling in 3D single spheroids^12,13^ remain limited due to the absence of a step-by-step protocol. Here we provide an optimized and detailed protocol using patient-derived brain tumor neurospheres. This protocol utilizes the Seahorse 3D spheroid assay kits and is specifically tailored for single spheroids derived from pediatric brain tumors, representing the first time this model has been systematically profiled using Seahorse technology in a 3D context.

### Cell culture

#### Timing: 3-4 days

Before starting the assay, make sure the cell line of interest is growing well in tissue culture. Timing refers to the time between passages. Seeding density, doubling times, and growth rates must be empirically determined for each cell line. Here we will provide specific instructions for the atypical teratoid rhabdoid tumor (ATRT) cell line, CHLA-05-ATRT.

**NOTE**: All cell culture steps, from preparation of media to passaging cells should be performed inside a biosafety cabinet using sterile techniques.

### Preparation of cell culture media

1. Pour two 500 mL bottles of DMEM/F12 through a 1 L, 0.45 µm filter, and add 10% Penicillin-Streptomycin. This constitutes the base media and should be stored at 4 °C.
2. Calculate the amount of base media needed.
3. Add the base media into a 50 mL conical tube.
4. To make working media, add the following to the base media: B-27 Supplement (Gibco, 17504001) at a final concentration of 2%, 20 ng/mL of EGF (Stem Cell Technologies, 78006.2), and 20 ng/mL of FGF (Stem Cell Technologies, 78003.2), and heparin solution 0.2% (Stem Cell Technologies, 07980) at a final concentration of 2 µg/mL.

**NOTE:** Prepare the working media fresh daily.

### Passage Cells

5. Collect the cells from an ultra-low attachment (ULA) T75 flask using a serological pipette and add the cells to a 50 mL conical tube.

6. If any cells are still attached to the flask, wash with 10 mL of sterile PBS and add this to the same 50 mL tube. Keep the ULA flask as this can be reused for up to two months.

7. Centrifuge the 50 mL conical tube at 340 x g for 4 minutes at 20 °C to pellet the cells.

8. Aspirate the supernatant without disturbing the cell pellet.

9. Resuspend the cell pellet in at least 1 mL of Accutase by pipetting up and down.

10. Centrifuge again at 340 x g for 4 minutes at 20 °C to pellet the cells.

11. Aspirate the supernatant without disturbing the cell pellet.

12. Resuspend the cell pellet in 1 mL of working media. Depending on the size of the pellet, add media up to 5 mL.

13. Add 10 µL of cell suspension to a 1.5 mL microcentrifuge tube and remove tube from the biosafety cabinet.

14. Add 10 µL of Trypan Blue to the same tube.

15. Add 10 µL of the cell suspension/Trypan Blue mixture to a Countess slide.

16. Using the Countess, record the percentage of live cells. live cell count, and total cell count.

17. Add media to a ULA flask and seed cells at the appropriate density depending on the cell line (the optimal seeding cell density for CHLA-05-ATRT was empirically determined to be 0.2 x 10^6^/mL).

## Key resources table

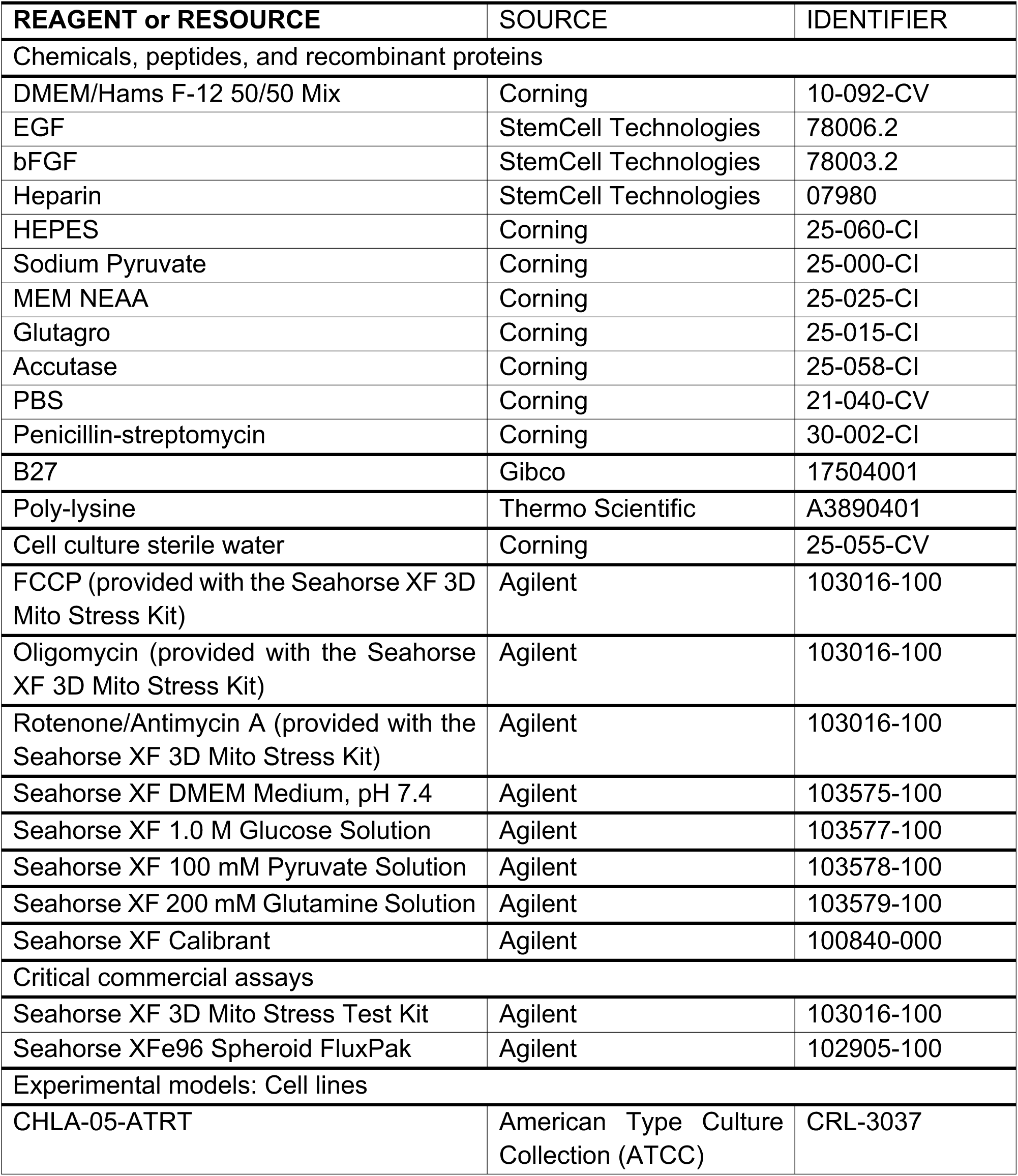
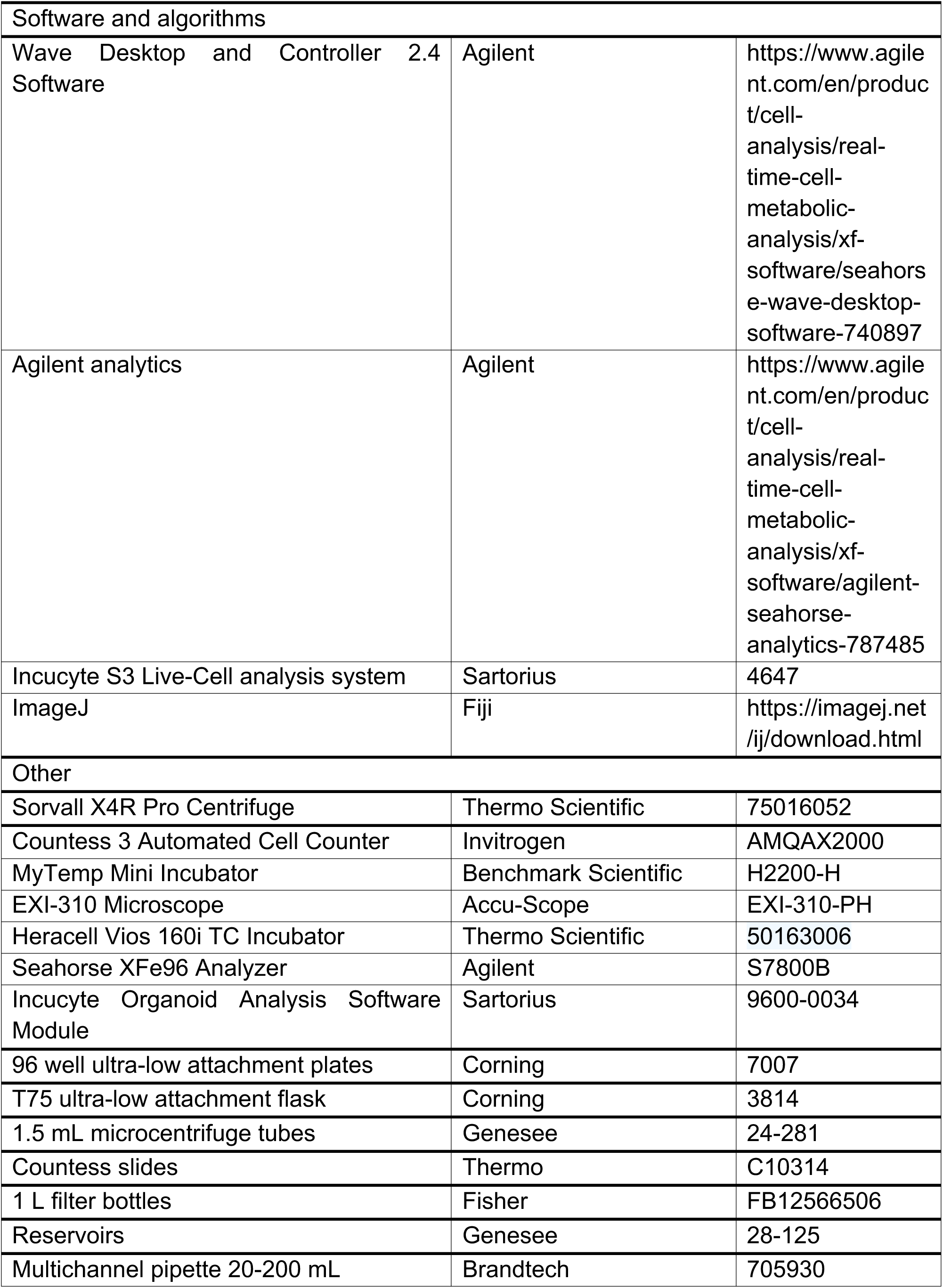
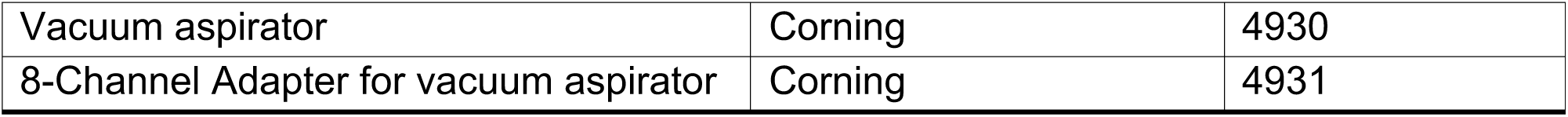

## Materials and equipment

### Agilent Seahorse XF analyzer

This protocol is optimized for the Seahorse XFe96 Analyzer instrument. General manufacturer’s instructions for 3D Spheroid FluxPak can be found at: https://www.agilent.com/cs/library/usermanuals/public/XFe96_Spheroid_Microplate_Users_Guide.pdf

General manufacturer’s instructions for the Seahorse XF Cell Mito Stress Test can be found at: https://www.agilent.com/cs/library/usermanuals/public/XF_Cell_Mito_Stress_Test_Kit_User_Guide.pdf.

Agilent provides other instruments that can be alternatively used, although they require optimization, including the Seahorse XF HS mini (8 wells) and the Seahorse XFe24 (24 wells) Analyzers.

**NOTE**: Although Agilent offers a Seahorse XF 3D Mito Stress Test Assay and protocol, this is designed for tissue samples and small organisms, and therefore it is not applicable for the approach used here.

### Preparation of assay medium

Here we use the Agilent Seahorse XF DMEM Medium, pH 7.4 without phenol red. In this protocol, we prepare the media following manufacturer’s instructions: Seahorse XF DMEM Medium is supplemented with 1 mM pyruvate, 2 mM glutamine, and 10mM of glucose (see the step-by-step section for more details). The assay media should be prepared fresh before the assay.

**NOTE:** Agilent provides alternative media that can be used with different cell lines. Medium composition can be adjusted based on the cell type and the experiment conditions.

## Step-by-step method details

This step-by-step protocol has been optimized for measuring cellular energetics in pediatric patient-derived brain tumor spheroid models. It includes the following sections: culturing neurosphere, poly-lysine coating of the Seahorse microplate, transfer of neurospheres to the pre-coated microplate, cartridge hydration, Seahorse XF 3D Mito Stress assay, and normalization methods.

### Culturing neurospheres

#### Timing: 1 hour for plating, 4-6 days for neurosphere formation

In this step we describe how to culture suspension cells to allow them to form neurospheres. All steps are performed in sterile conditions.

The number of cells per well depends on the cell line and the incubation period between plating and the assay. This step should be empirically determined to ensure that neurospheres reach an appropriate size by the time they are transferred to the microplate. Here, we seeded 5000 cells per well in a 96-well ULA plate and let them grow for 4-6 days.

**NOTE**: Plating neurospheres at the edge of the plate is not recommended since the plate will be incubated for several days and evaporation will occur. Add an equal volume of sterile PBS to each of the outer wells of the plate. By doing this you will have only 60 wells available for neurosphere plating.

1. Follow steps 1 to 12 from the passage cells section above.

2. Calculate the volume of cells needed for a 96-well ULA plate. An example is shown below.

a. Add the calculated volume of media to a 15 ml tube.
b. Subtract the cell volume.
c. Add the cell volume into the media.
d. Gently pipet up and down to evenly distribute the cells in the media.

Example:

Cell count: 1 x 10^5^/mL

Desired cell density: 5000 cells in 100 µL per well for a total of 96 wells

Total number of cells: 5000 x 96 = 48,000 cells

Total volume of cells to collect: 48,000/1 x 10^5^ = 480 µL

Total volume of media needed:100 x 96 = 9600 µL = 9.6 mL

Therefore: Collect 480 µL of cells in 9.120 mL of media

3. Add 100 µL of sterile PBS to the outer wells of a 96-well ULA plate using a reservoir and a multichannel pipette.

4. Plate 100 µL of the cells and media prepared above in each well of the 96 well plate using a reservoir and a multichannel pipette.

a. Once all cells have been plated, spin the plate at 200 x g for 15 minutes at 20 °C.
b. Check the plate under a microscope to ensure the cells have come together at the center of the wells.

5. Incubate the plate in an incubator at 37 °C with 5% CO_2_.

6. Let the neurospheres growth for 4-6 days (**Figure 1**)

**Figure 1.**
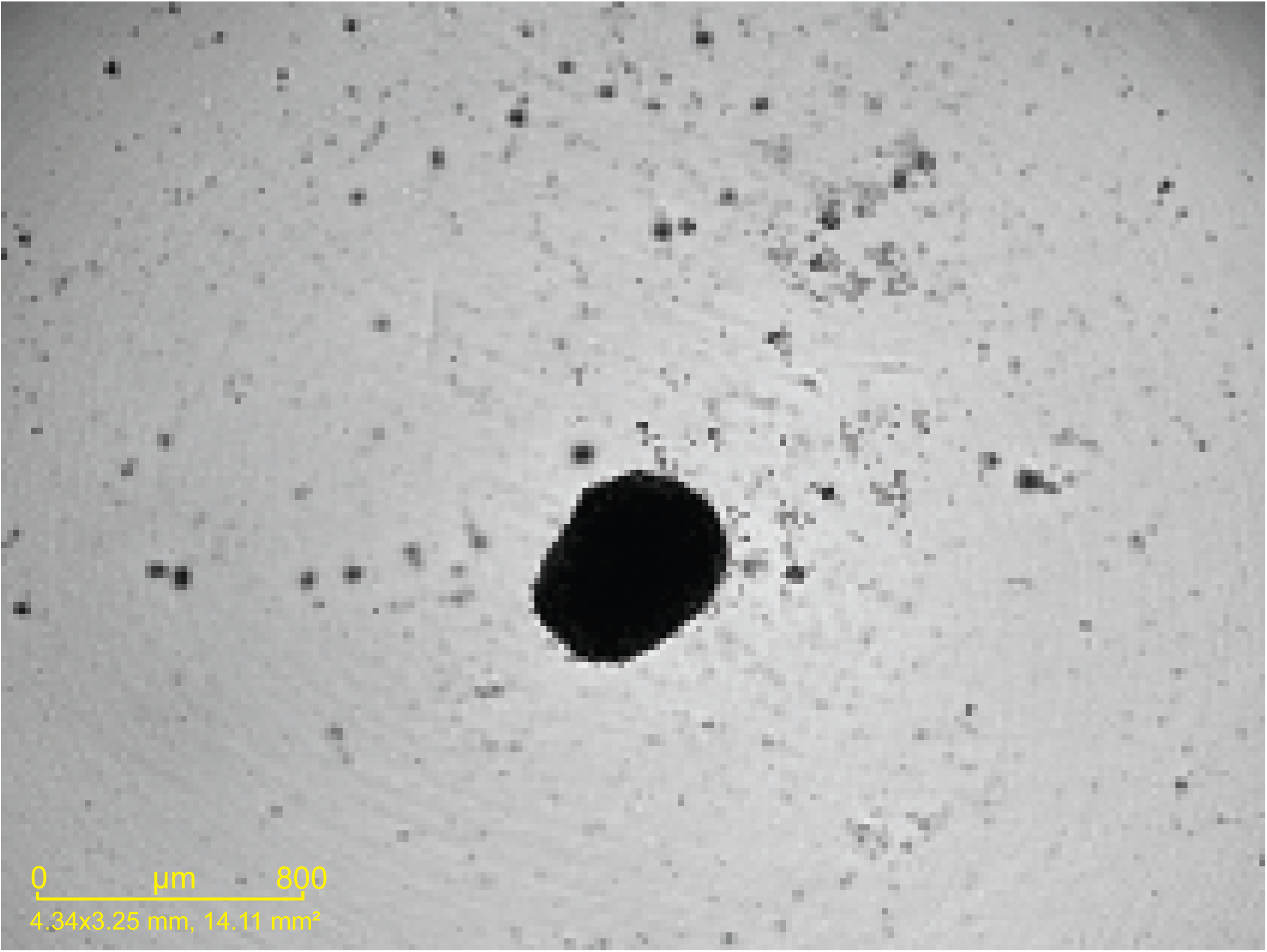
Image of spheroid taken 5 days after plating (5000 cells were seeded).

**NOTE:** When possible, prepare a larger volume of cells than what is needed.

### Poly-lysine coating of assay microplate

#### Timing: 3-4 hours

Neurospheres aggregate with each other but do not attach to the surface of the tissue culture vessels. To promote cell adhesion, poly-lysine is used to coat the wells of the assay microplate. This step is critical to avoid spheroid movement during the assay, which would compromise the accuracy and reliability of the results (Troubleshooting).

7. Add 50 µL of poly-lysine to each well of the assay microplate, using a multichannel pipette.

a. Cover the plate and let it sit for one hour inside the biosafety cabinet.
b. Aspirate the poly-lysine from each well, using a vacuum aspirator. We recommend using a multichannel aspirator for convenience; however, a single aspirator may also be used.
c. Wash the plate three times with sterile water. **NOTE:** This step is critical as excess poly-lysine can be toxic to the cells.
d. Keep the plate uncovered inside the biosafety hood and let it sit to dry for two hours or as long as needed.

8. Use the plate immediately for neurosphere transfer.

a. If not used immediately, seal the plate with Parafilm and store it at 4 °C. The plate should be used within one week.

### Transferring neurosphere to the coated microplate

#### Timing: 45 minutes – 1.5 hour (variable depending on how many spheroids need to be transferred)

Here, we describe how to retrieve spheroids from the culture plate and transfer them to the pre-coated assay microplate, once the spheroids reach the desired size. While this transfer process can be tedious, the following tips can help ensure an efficient and successful transfer process.

NOTE: This step should be completed the day before the assay

NOTE: We recommend practicing the transfer process as much as possible before running the first assay to reduce transfer time and ensure consistency.

**NOTE**: It is critical to visualize the spheroid inside the plate at all times.

9. Add 20 µL of culture media to the poly-lysine coated microplate.

This facilitates the release of the spheroid from the pipette tip. Do not add a larger volume of media, as this can cause the spheroids to float around and prevent them from attaching to the center of the well.

10. Carefully remove as much media as possible from each culture plate well without disturbing the spheroids.

a. This helps with the spheroid retrieval.

11. Set the pipette to a maximum of 20 µL.

a. This helps minimize excessive media during spheroid retrieval.

12. Using a 200 µL pipette and 200 µL tips, place the tip straight (perpendicular) against the bottom of the well (**Figure 2**).

**Figure 2.**
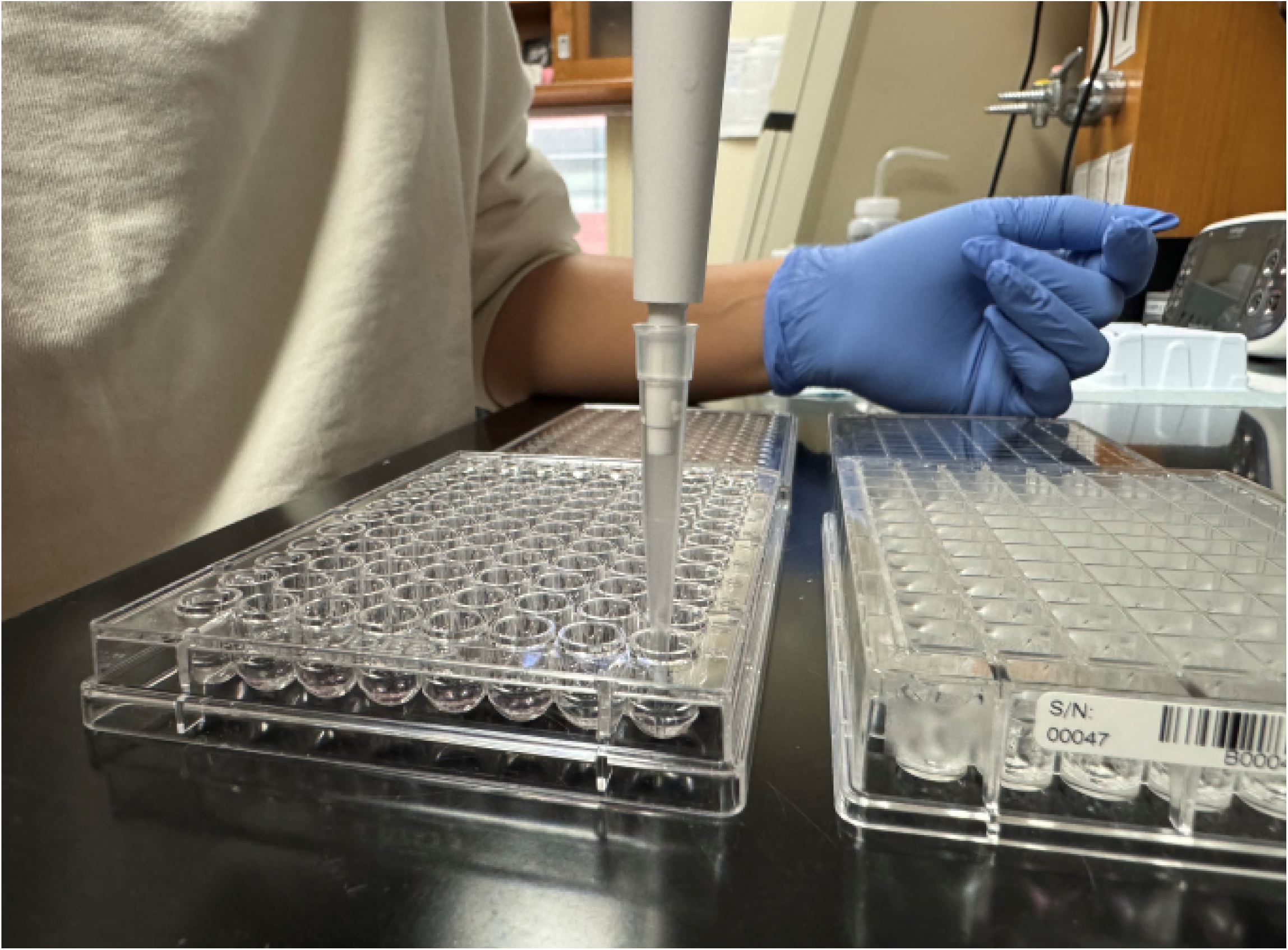
Image showing the positioning of the tip against the bottom of the 96-well ULA plate.

13. Gently aspirate single spheroids by picking up to 20 µL of media.

**CRITICAL**: Avoid introducing air bubbles inside the pipette tip. If needed, hold the pipette without releasing the plunger to prevent air intake. Ideally, position the spheroid at the tip of the pipette for easy transfer and to minimize disruption (**Figure 3**).

**Figure 3.**
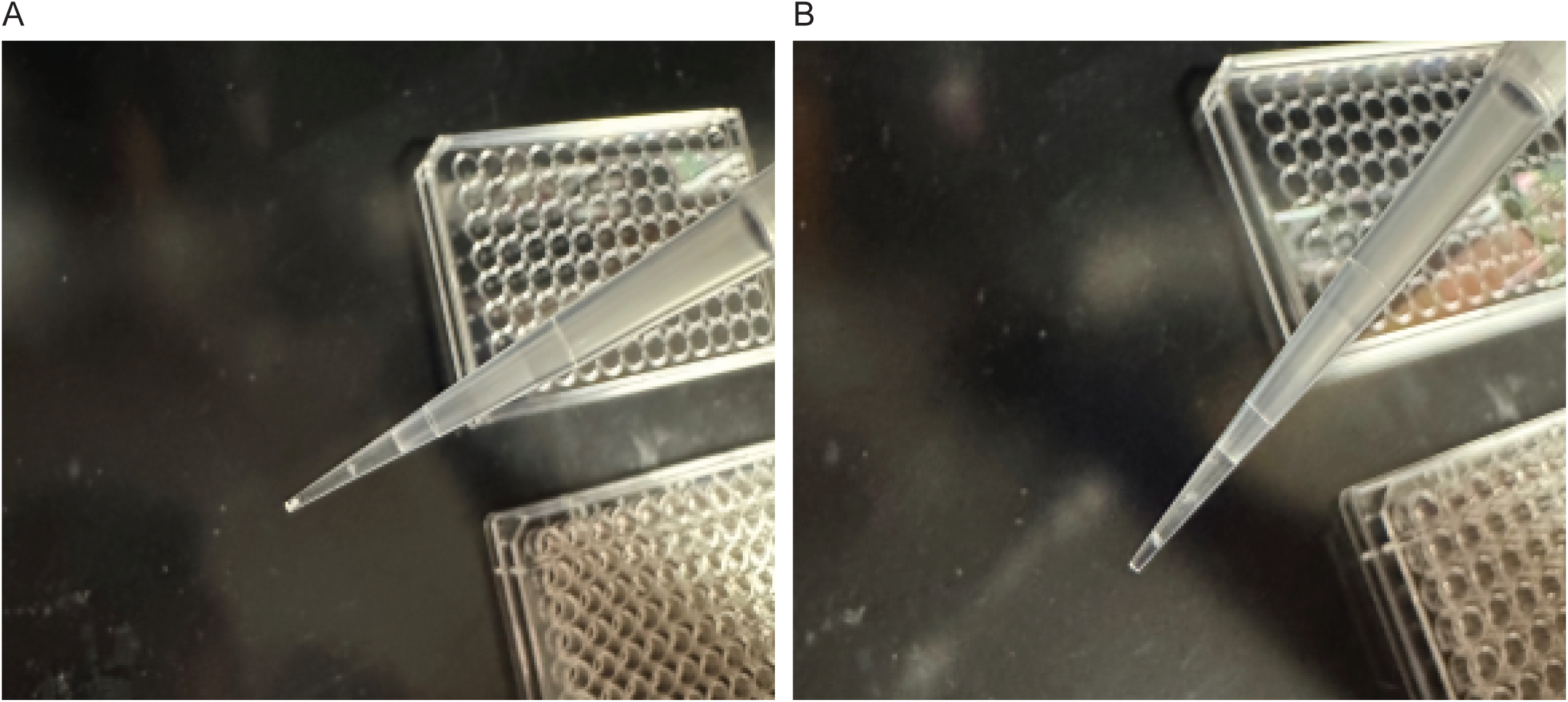
Images showing the spheroid at the very tip **(A)** of the pipette or very close to the tip **(B)**.

14. Place the tip against the bottom center of the microplate well. Hold steady and gently release the spheroid to allow it to settle into the well (**Figure 4**).

**Figure 4.**
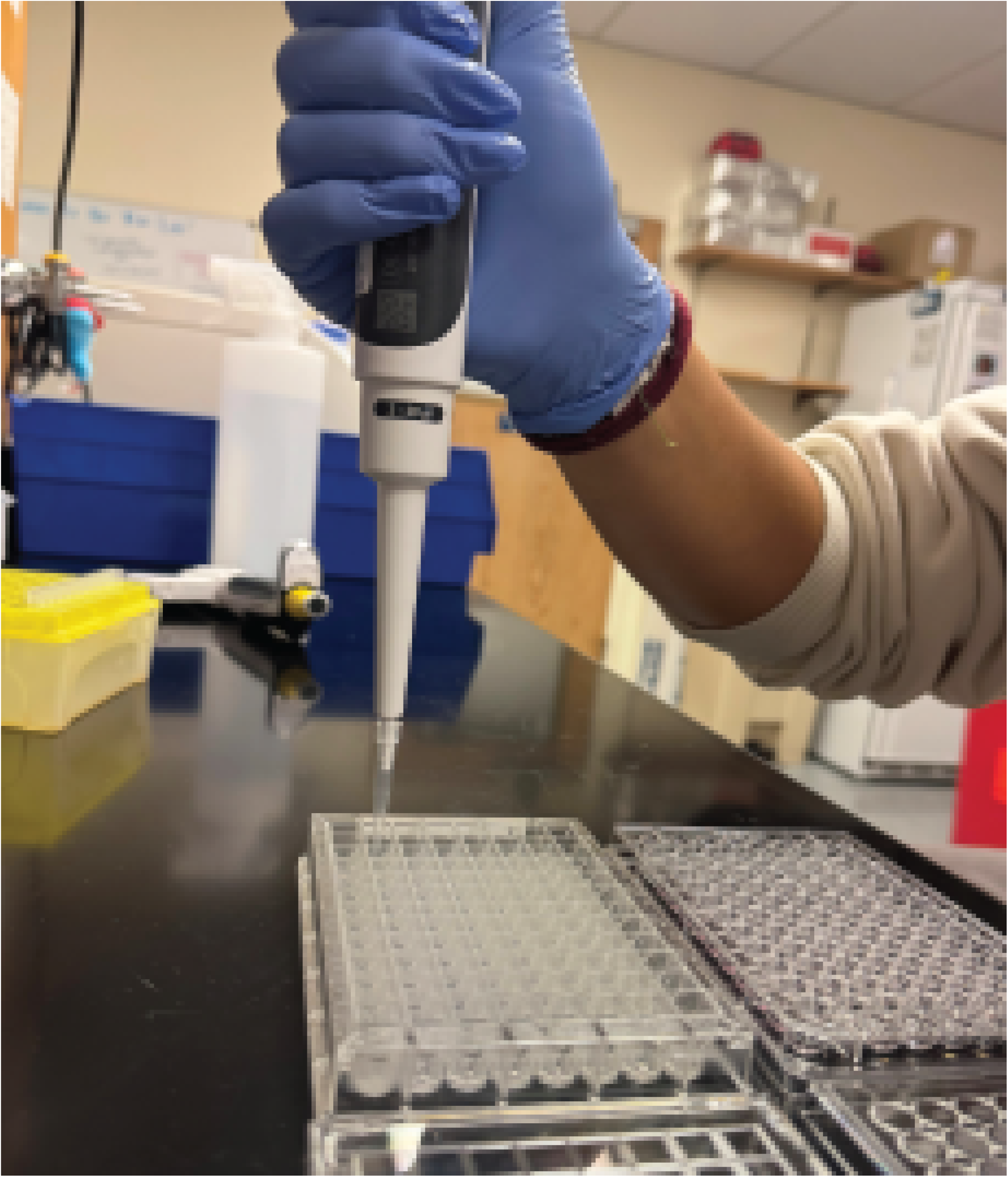
Image showing the position of the tip against the bottom of the assay microplate.

**CRITICAL**: Do not add spheroids to the four corner wells, as these will be used by the software for background normalization.

**CRITICAL**: Do not pipette too forcefully as this may create bubbles and cause disruption of the spheroid.

15. Slowly add 100 µL of culture media using a multichannel pipette.

16. Visually confirm that the spheroids are present and centered within each well (**Figure 5**).

**Figure 5.**
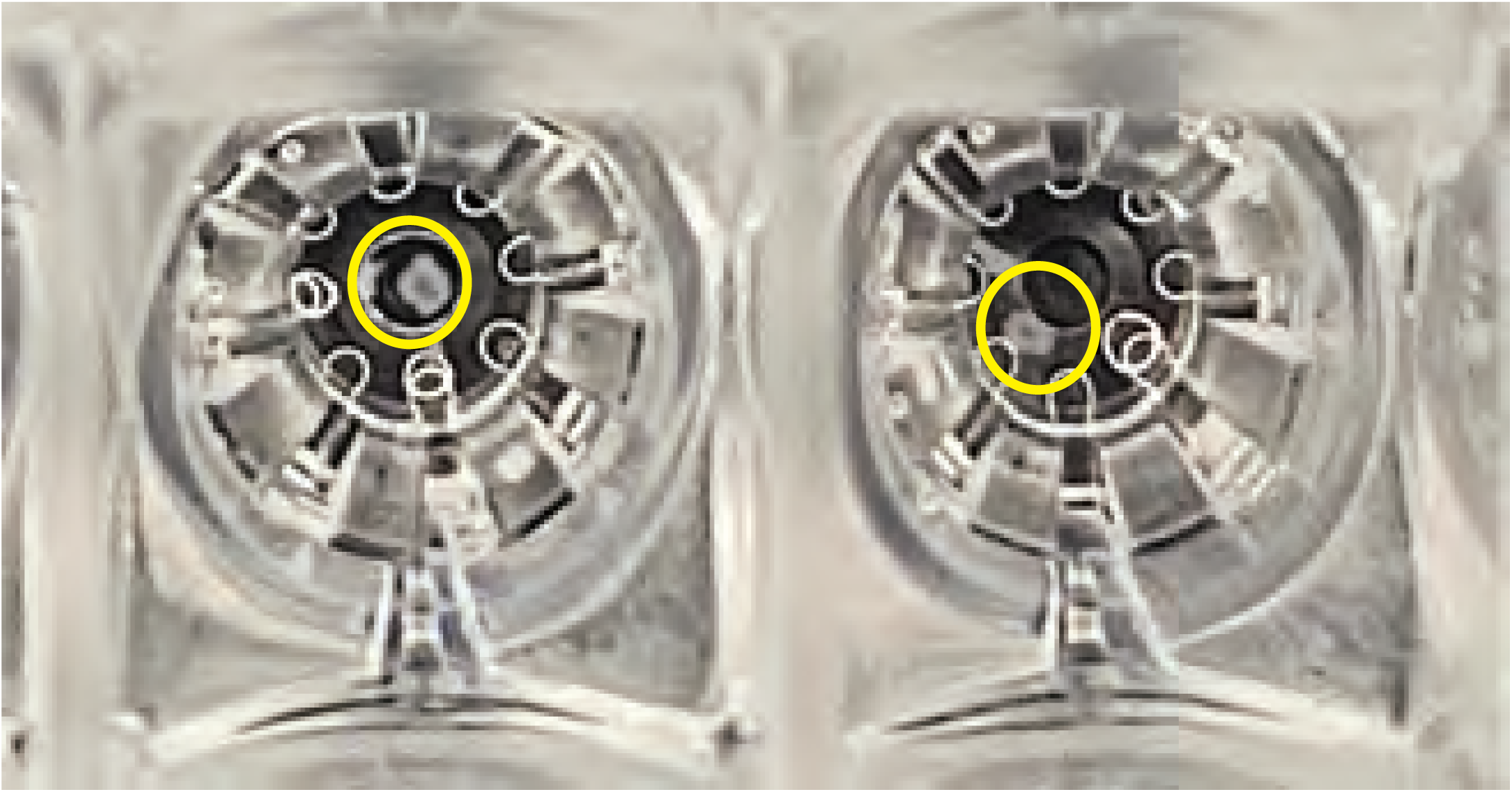
Image of two selected wells of the assay microplate. The yellow circle highlights the spheroids within the wells. On the left, the spheroid is centered, on the right the spheroid is at the edge of the well.

17. Place the microplate in an incubator at 37 °C with 5% CO_2._

**NOTE**: The plate was opened and manipulated outside the biosafety cabinet and is therefore no longer sterile. We utilize a different 37 °C, 5% CO_2_ incubator than the one for tissue culture.

**NOTE:** Take pictures of the microplate before and after the assay to ensure presence of spheroids and for normalization at the end of the assay.

### Cartridge hydration

#### Timing: Overnight

**CRITICAL**: This step should be completed the day before the assay

Cartridge hydration must be performed one day prior to conducting the assay. The Seahorse cartridge is an essential component as it presents sensors that detect changes in pH and oxygen concentration. The sensors must be thoroughly hydrated to function properly.

For this step we followed manufacturer’s instructions that can be found here: https://www.agilent.com/cs/library/usermanuals/public/user-manual-agilent-seahorse-xf-hydrobooster-5994-4361en-agilent.pdf

18. Remove the contents of the Seahorse XFe96/XF Pro Extracellular Flux assay kit, which include the XFe96/XF Pro Sensor Cartridge (green), Hydrobooster (pink), and Utility Plate (clear) (**Figure 6**).

**Figure 6.**
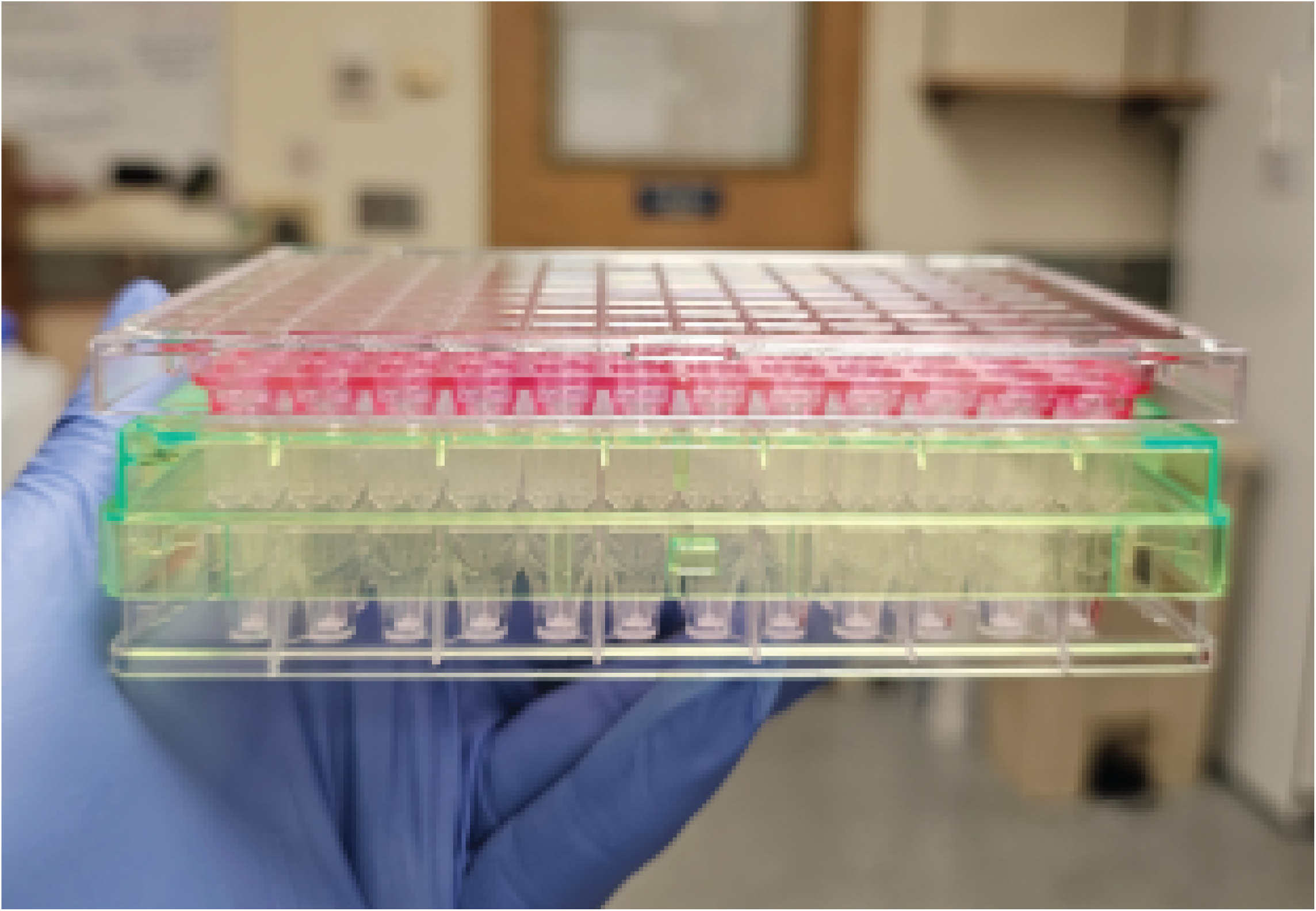
Seahorse XF Pro Sensor Cartridge (green), Hydrobooster (pink), and Utility Plate (clear).

19. Remove the sensor cartridge and place it upside down (the sensors at the bottom of the cartridge should not touch any surface).

20. Using a multichannel pipette, add 200 µL XF Calibrant to each well of the utility plate.

21. Place the XF Hydrobooster on top of the utility plate and push down to ensure it tightly covers the utility plate.

22. Place the sensor cartridge on top of the Hydrobooster into the opening of the utility plate.

23. Verify that the sensors are submerged in the calibrant.

24. Place the cartridge with the Hydrobooster and the utility plate in a non-CO_2_ incubator at 37 °C and leave it overnight.

**NOTE**: The utility plate is only used to hydrate the cartridge, and it will be replaced by the spheroid-containing microplate before the assay begins.

**CRITICAL**: Evaporation of the XF Calibrant can compromise the hydration steps and ultimately, the accuracy of the data. To prevent this, place a tray of water inside the incubator to maintain a humidified environment. Skipping this step will result in calibrant evaporation.

25. Turn on the Agilent Seahorse XFe96 Analyzer and let it warm up overnight. This step is critical to allow equilibration of the thermal tray.

### Preparation of assay medium

#### Timing: 30 minutes

This step must be performed under sterile conditions within a biosafety cabinet before starting the assay, as the media needs to be warm.

26. Prepare assay medium by adding the following reagents to a 50 mL conical tube.

|  |  |
| --- | --- |
| Seahorse XF DMEM Medium, pH 7.4 | 48.5 mL |
| Seahorse XF 1.0 M Glucose Solution | 500 μL |
| Seahorse XF 100 mM Pyruvate Solution | 500 μL |
| Seahorse XF 200 mM Glutamine Solution | 500 μL |

27. Warm the assay medium to 37 °C in a water bath and keep warm until use.

### Seahorse XF 3D Mito Stress assay

#### Timing: 2-3 hours

28. Take the microplate with cells out of the incubator and prepare it for the assay.

29. Remove the culture media from each well without disturbing the spheroids. We recommend using a multichannel pipette and not an aspirator.

**NOTE**: All media should be removed. Use a single channel pipette to remove excess media from each individual well, if needed.

30. Gently add 175 µL of the warm assay medium to each well using a multichannel pipette, including the four corner wells that do not contain spheroids.

31. Incubate the plate in an incubator without CO_2_ at 37 °C for 1 hour.

**CRITICAL**: Do not exceed the 1 hour incubation time.

32. Prepare the drug stocks by resuspending them in the assay medium as follows:

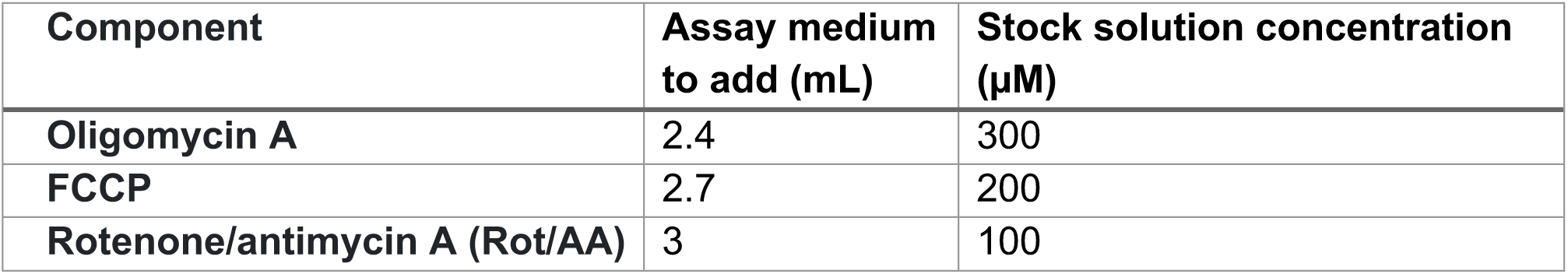

| Component | Assay medium to add (mL) | Stock solution concentration (μM) |
| --- | --- | --- |
| Oligomycin A | 2.4 | 300 |
| FCCP | 2.7 | 200 |
| Rotenone/antimycin A (Rot/AA) | 3 | 100 |

33. Vortex briefly to dissolve completely the contents of the drug tubes.

34. Prepare the drug working solutions based on the final concentration to be injected into the ports. In this protocol, we prepared the working drug solutions as follows:

| Component | Assay medium to add (mL) | Stock solution to add (μL) | Working concentration (10X) (μM) |
| --- | --- | --- | --- |
| Oligomycin A | 2.75 | 250 | 25 |
| FCCP | 2.25 | 750 | 50 |
| Rotenone/antimycin A (Rot/AA) | 2.85 | 150 | 5 |

35. For each drug working solution, load the amount indicated in the table below into the respective loading ports (Oligomycin A: Port A, FCCP: Port B, Rotenone/antimycin A: Port C – **Figure 7**) of the cartridge. Port D is left empty. Use a multichannel pipette to minimize pipetting errors. Prepare an excess amount of drug working solutions to ensure sufficient volume. Use the microplate guide provided within the kit to help load the drugs.

a. Add the drugs to the four corner wells that do not contain spheroids.

**Figure 7.**
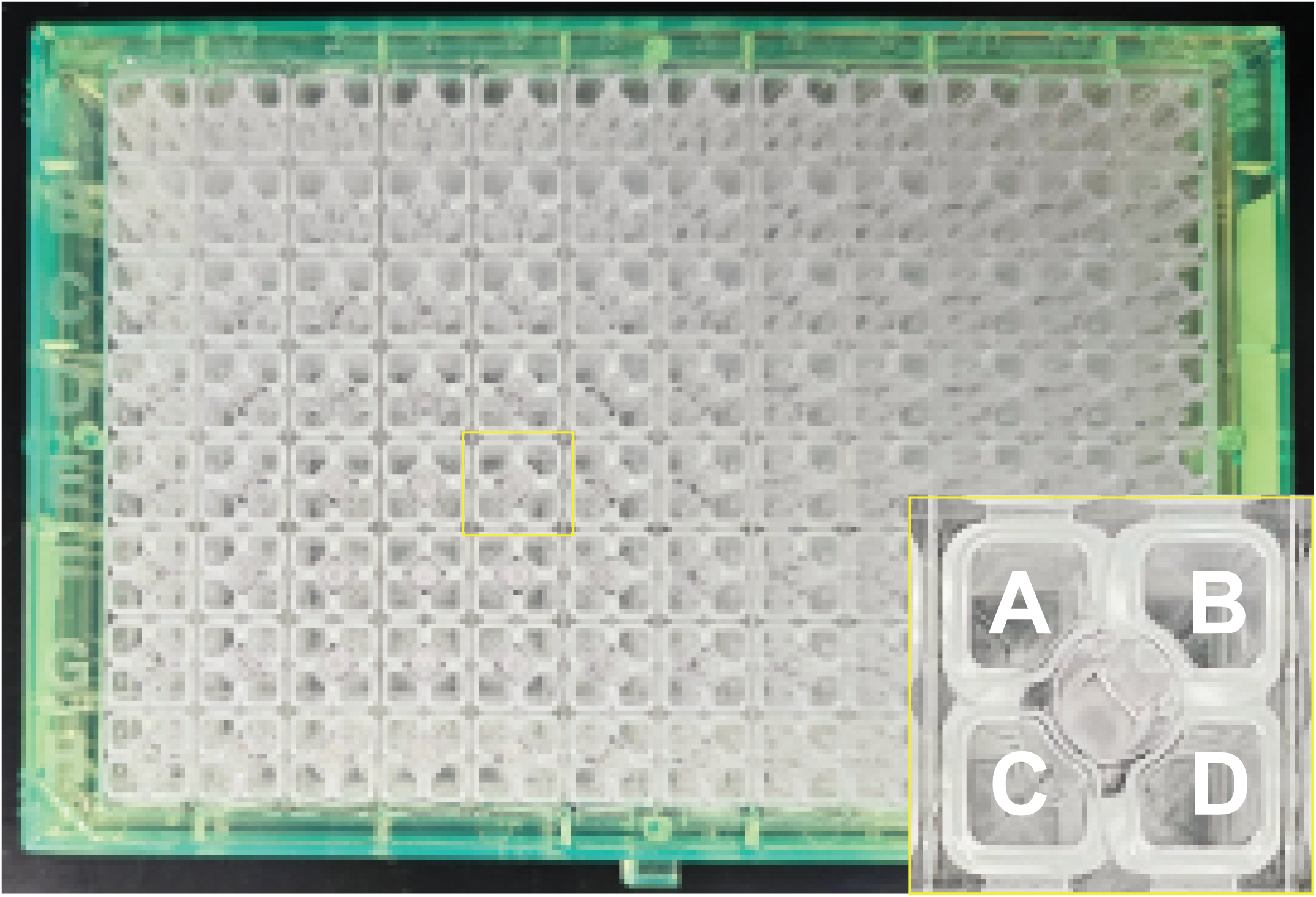
Image of sensor cartridge. The yellow box highlights the well loading ports (A, B, C, D).

| Component | Port | Volume added to port (μL) | Final concentration per well (μM) |
| --- | --- | --- | --- |
| Oligomycin A | A | 20 | 2.5 |
| FCCP | B | 22 | 5 |
| Rotenone/antimycin A (Rot/AA) | C | 25 | 0.5 |

**NOTE**: Final FCCP concentration per well should be optimized based on specific cell type and experimental conditions (Troubleshoot).

**NOTE:** Following cartridge preparation, the cartridge will be placed inside the XF Analyzer as prompted by the software for calibration test. This can take 10-15 minutes. If 1 hour is not sufficient to complete all these steps, prepare the stock and drugs solution prior to the 1 hour incubation of the microplate.

36. Open the Seahorse XFe96 Analyzer Wave Software.

37. Open the XF Cell Mito Stress test assay file and fill in the assay details. You may include details about the cell lines and media. Be sure to make a plate map with the group conditions. Before starting the assay, ensure the protocol default setting is edited as needed for your experiment. The protocol settings used in this protocol are shown in Table 1.

**Table 1.** XF Cell Mito Stress Test Assay Setting.

|  |  |  |  |
| --- | --- | --- | --- |
| **Basal** | **3 cycles** | **Mix** | **3 min** |
|  |  | Wait | 0.40 min |
|  |  | Measure | 3 min |
| **Oligomycin** | 5 cycles | Mix | 3 min |
|  |  | Wait | 0.40 min |
|  |  | Measure | 3 min |
| **FCCP** | 5 cycles | Mix | 3 min |
|  |  | Wait | 0.40 min |
|  |  | Measure | 3 min |
| **Rotenone + Antimycin A** | 10 cycles | Mix | 3 min |
|  |  | Wait | 0.40 min |
|  |  | Measure | 3 min |

38. When ready to begin the assay, click Start Run.

39. When prompted by the software, place the cartridge with the utility plate in the instrument tray and follow the software instructions. Make sure to remove the Hydrobooster and lid before loading the cartridge. The software will perform equilibration and calibration, which take approximately 15 minutes.

40. At the end of the calibration, remove the utility plate (the cartridge will remain inside the instrument automatically) and load the assay microplate as prompted by the software. Remove the lid before loading the plate.

41. At this point, the assay will begin automatically. Oxygen consumption rate (OCR) and Extracellular Acidification Rate (ECAR) measurements will be displayed in real time on the Wave Software screen, enabling an initial inspection of the results. Examples of a successful OCR measurement profile when the spheroid is visible in the well, compared to a failed OCR profile in the absence of a spheroid are shown in **Figure 8**.

**Figure 8.**
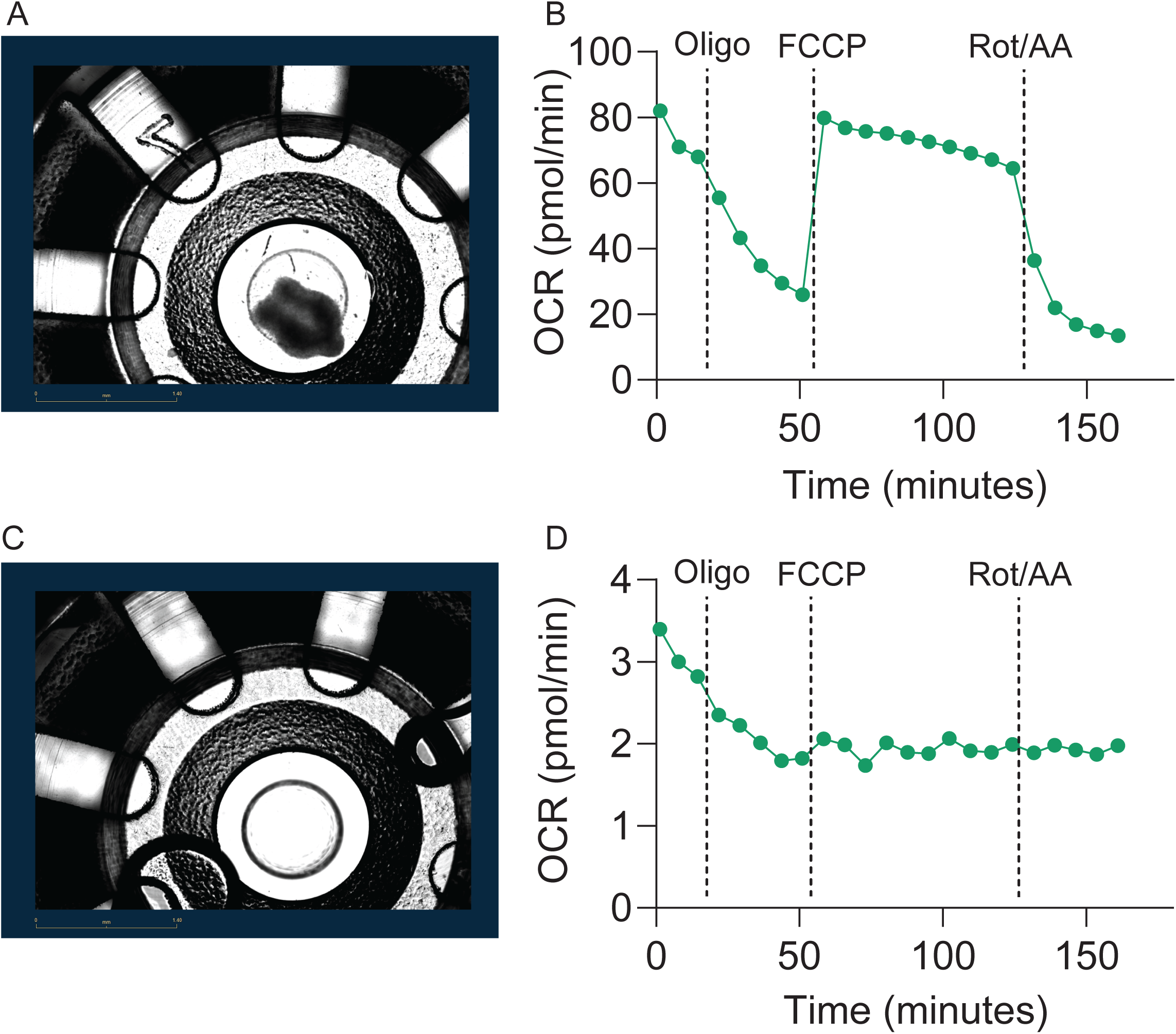
**(A)** Picture of spheroid in the center of the microplate well at the end of the assay and **(B)** the respective OCR kinetic graph showing a response after drug injections as expected. **(C)** Picture of a microplate well with no spheroid detected at the end of the assay and **(D)** respective OCR kinetic graph showing no response after drug injections.

**NOTE:** The optimal number of cycles must be determined empirically. The response of 3D spheroids to drug injections differs from that of monolayer cells and typically requires longer time to reach the full effect. For the Mito Stress assay, the default setting is three cycles after drug injection. Ensure that a stable response of at least three measurements is obtained. We tested up to ten measurement cycles for each drug and observed that response reached a plateau after five cycles for Oligomycin and FCCP, while Rotenone/AA requires ten cycles to result in complete inhibition.

### Normalization

#### Timing: 30 minutes-1.5 hour (variable depending on how many wells need to be quantified)

The bioenergetic parameters can vary due to differences in spheroid size. Spheroid size can change due to specific treatments, genetic manipulation that affects proliferation, or even technical variability during the seeding or the transfer from the culturing plate to the assay microplate. Therefore, normalization of the bioenergetics parameters to the number of cells present in each well at the time of the assay is essential. We have determined that measuring the spheroid area at the end of the assay provides the most accurate and reliable normalization method.

41. At the end of the assay gently remove the microplate and the cartridge from the instrument.

42. Discard the cartridge and put the microplate inside the Incucyte.

43. Open the Incucyte software.

44. Image the plate once following the instructions prompted in the software, using the spheroid analysis application.

45. Once the imaging is complete, save the picture and open it with ImageJ.

a. Select the Polygon Selection.
b. Draw on the spheroid area being as precise as possible (**Figure 9**).
c. Click Analyze and then Measure (or Ctrl+M).
d. This will automatically open a window with all the parameters measured.
e. Copy it and paste it to an Excel spreadsheet.

**Figure 9.**
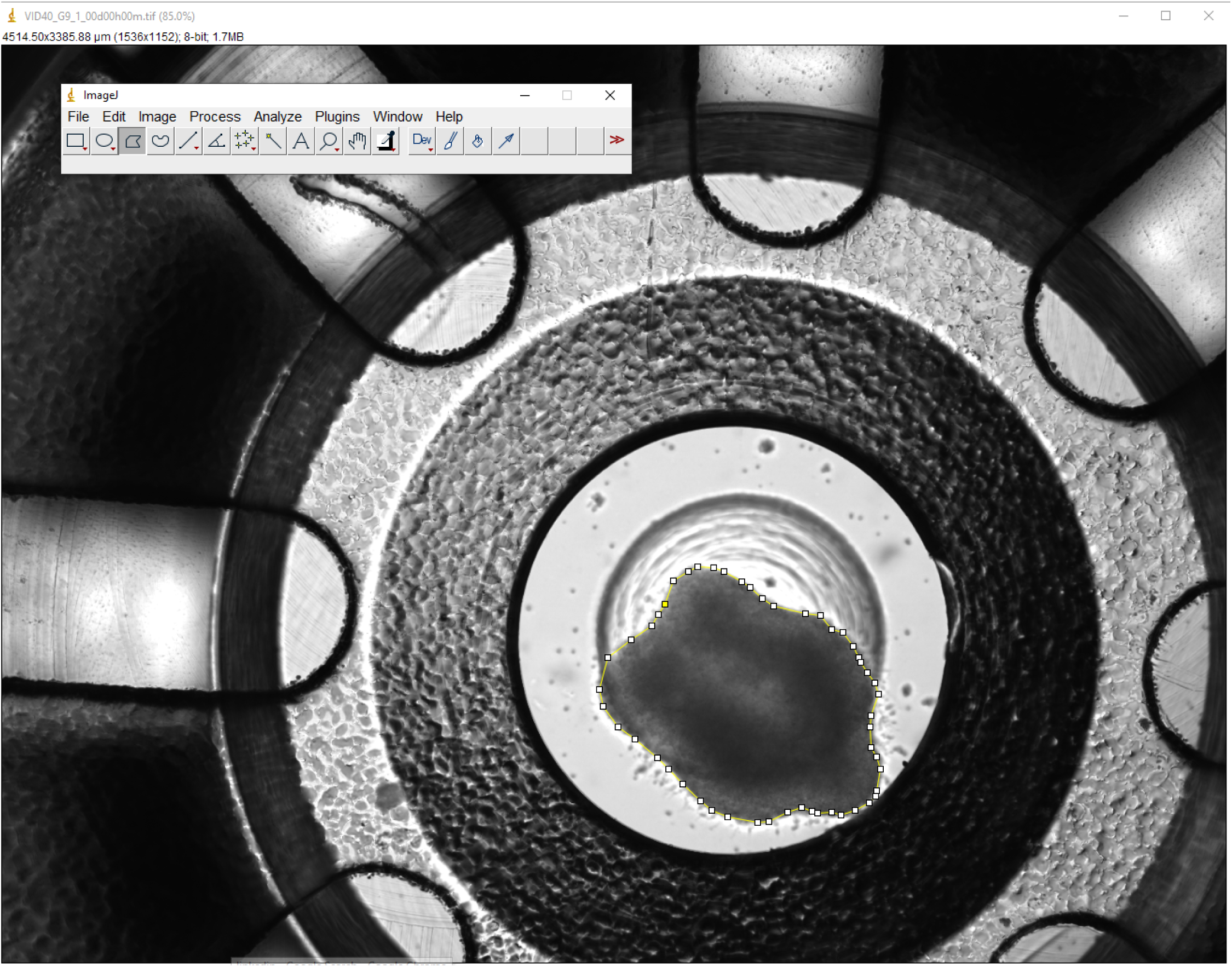
The spheroid area is marked (yellow line and white dots) using the polygon selection for analysis in ImageJ.

46. Open the Mito Stress assay result data in Agilent Wave software.

47. Click on Normalize and add each area value to the corresponding well.

48. Raw data can be exported as an Excel file or a GraphPad Prism file.

**NOTE:** Make sure that the normalization option (slider bar) is on (blue) before downloading the raw data.

**NOTE**: We always take images of the microplate before and after the assay to ensure the spheroids maintain their size and structure throughout the experiment.

Other normalization approaches we tested included collecting spheroids from each well of an additional culture prepared and maintained under the same conditions at the assay plate. However, counting from a small number of cells can be inaccurate and unreliable. In addition, although we expect these spheroids to be the same as the ones transferred to the assay microplate, these spheroids were not used for the assay. This method is not sufficiently precise or direct.

We also attempted to use the Incucyte imaging and spheroid analysis tools for automated area measurement. However, the software is not currently optimized for imaging and analyzing spheroids in the Seahorse 3D 96-well microplate format, making this approach of limited utility.

DNA quantification is another potential normalization method, but it requires optimization. The DNA yield from small individual spheroids can be low and inaccurate. Moreover, collecting spheroids directly from the assay microplate at the end of the assay for DNA isolation may be technically challenging.

## Limitations

This protocol was optimized in the CHLA-05-ATRT cell line. This particular cell line forms spheroids that maintain a relatively round shape and remain compact throughout the assay, making them easier to handle and transfer from one plate to another. We also used a different cell line, CHLA-06-ATRT, that forms less compact spheroids. Although we were able to complete the Mito Stress assay in this cell line, we encountered challenges in consistently transferring the spheroids to the microplate. Transferring spheroids from the culture plate to the microplate can disrupt their structure, especially when the spheroids are loose, making downstream quantification and analysis more challenging. In addition, measuring the area does not account for cell density – some spheroids may appear larger simply because the cells are loosely aggregated, therefore size alone may not always accurately reflect the actual number of cells in a single spheroid.

These considerations are important when designing experiments with specific cell lines. Performing optimization for each cell line and experimental conditions before the full experiment is critical.

### Troubleshooting

Problem 1

The spheroids do not respond to the drug injections because they move from the center of the well or are disrupted during the assay.

Potential solution

Neurospheres do not adhere to the well of the microplate, therefore pre-coating the microplate is critical. It is important to optimize the coating based on your specific cell type and experimental conditions. We tested Geltrex and poly-lysine coating. We found that at the end of the assay, the spheroids were displaced in the Geltrex-coated wells. In contrast, the spheroids were attached to the poly-lysine-coated microplate wells only when spheroids were transferred the day before and allowed to settle overnight. When spheroids were transferred to the poly-lysine-coated wells on the same day as the assay, they were displaced by the end of the experiment **(Figure 10**).

**Figure 10.**
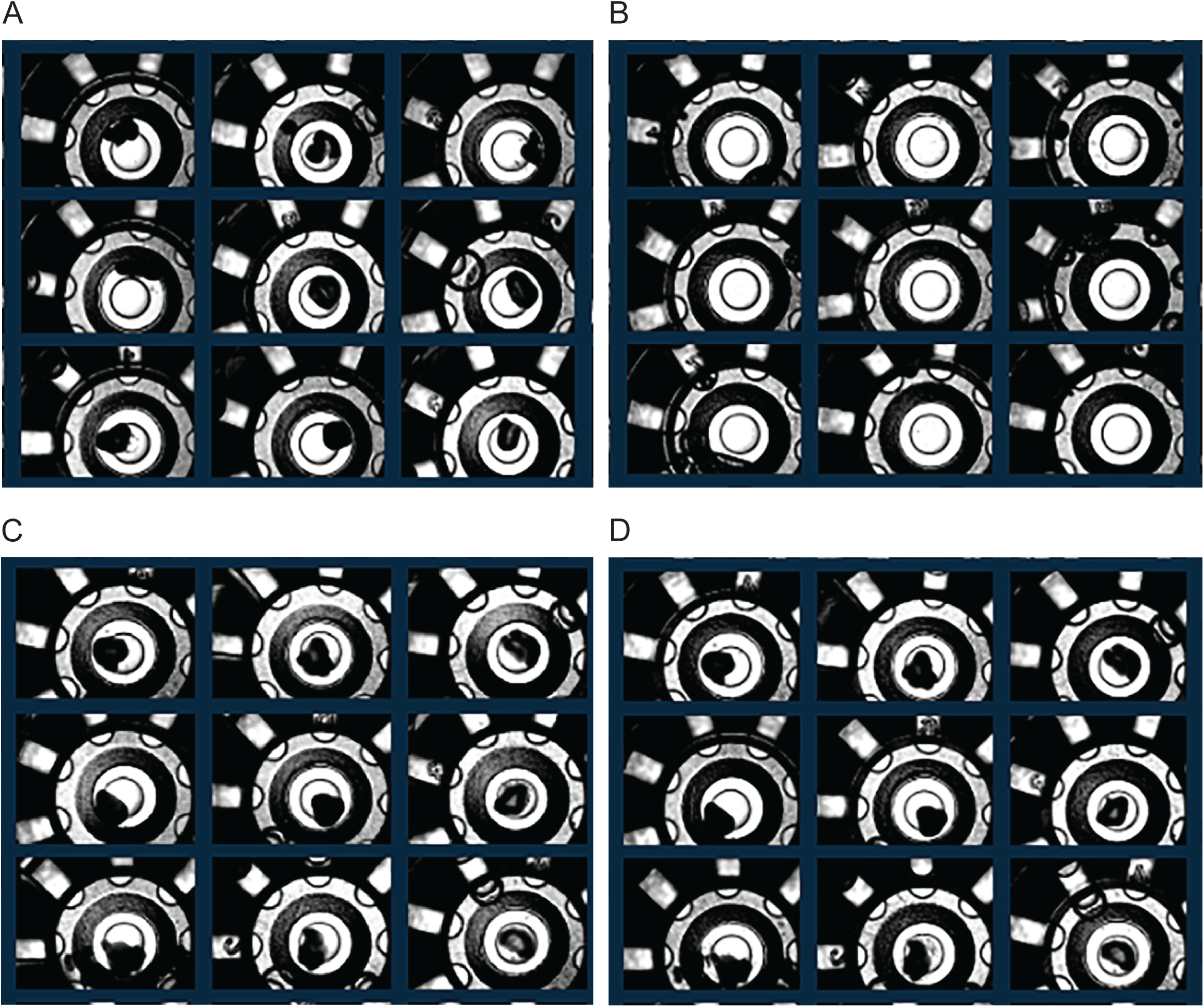
**(A)** Image of a poly-lysine coated microplate with spheroids transferred on the same day as the assay, taken before the assay. **(B)** Image of the same plate taken after the assay, showing that the spheroids were displaced and/or destroyed during the assay. **(C)** Image of a poly-lysine coated microplate with spheroids transferred the day before the assay, taken before the assay and **(D)** after the assay, showing that the spheroids were intact.

Problem 2

The cells respond to Oligomycin but do not respond to FCCP injection.

Final FCCP concentration per well should be optimized based on specific cell type and experimental conditions. The lowest concentration that causes maximal response should be selected. We tested 2 µM, 5 µM, and 10 µM FCCP and determined that only 5 µM provided an optimal response **(Figure 11)**.

**Figure 11.**
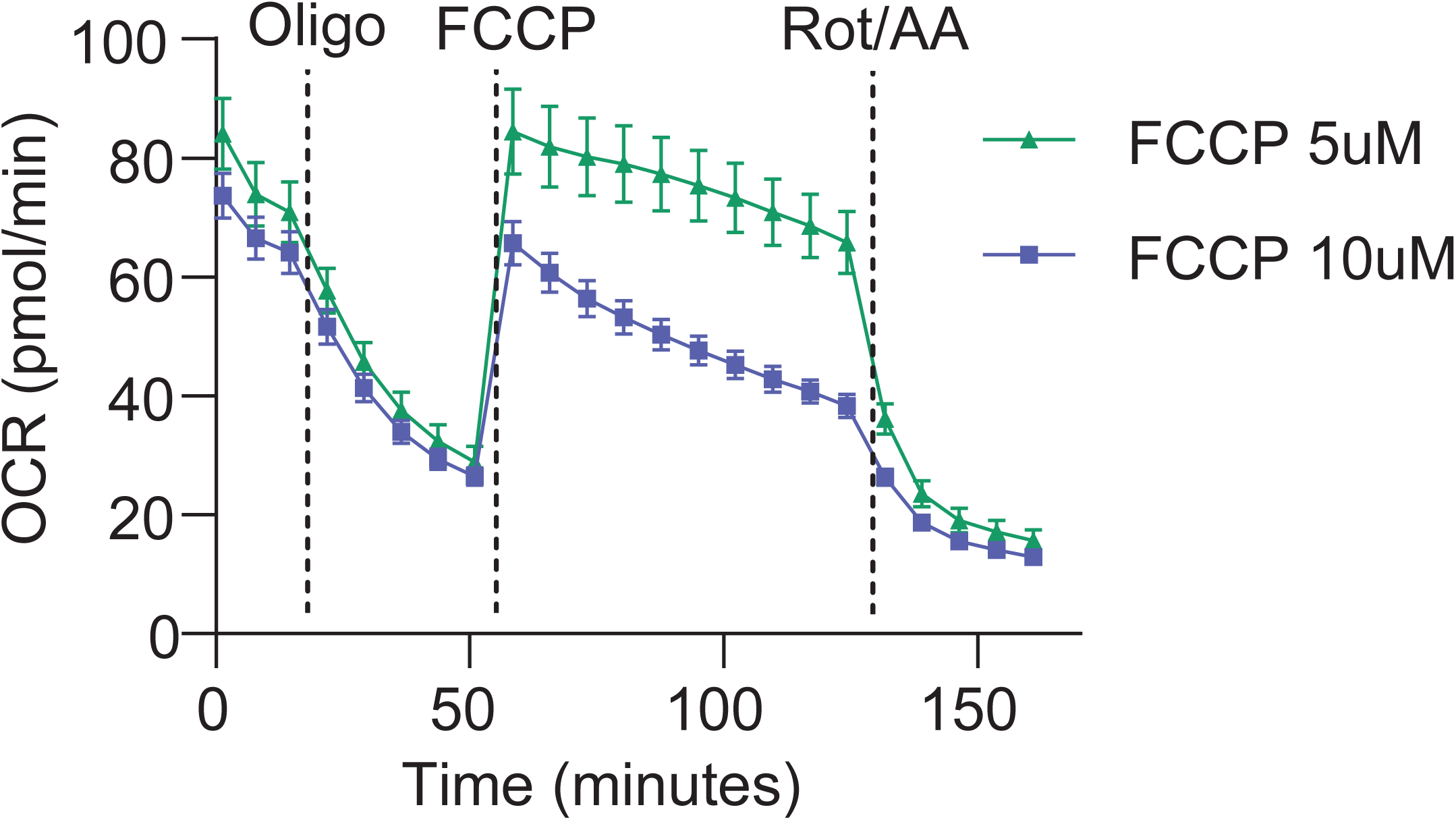
OCR graph shows the kinetic response after 5 µM of FCCP (green) and 10 µM of FCCP (blue).

## Expected outcomes

In this protocol, we optimize the Seahorse XF 3D Mito Stress Assay in single spheroids derived from pediatric brain tumors. Here we provide an example of the assay performed in neurospheres where CRISPR/Cas9 knockout of a candidate gene was performed, using two different guide RNAs (sgRNAs), sg2 and sg4 in **Figure 12**. Knockout was performed prior to the assay using lentiviral infection with the cells growing in suspension. After two days of antibiotic selection, the cells were plated for neurosphere formation for six days. The knockout resulted in a significant reduction in both basal and maximal mitochondria respiration (**Figure 12**). Given the effect of the knockout on cell viability, we measured the spheroid area using ImageJ and normalized the OCR results to the spheroid area using Seahorse Analytics.

**Figure 12.**
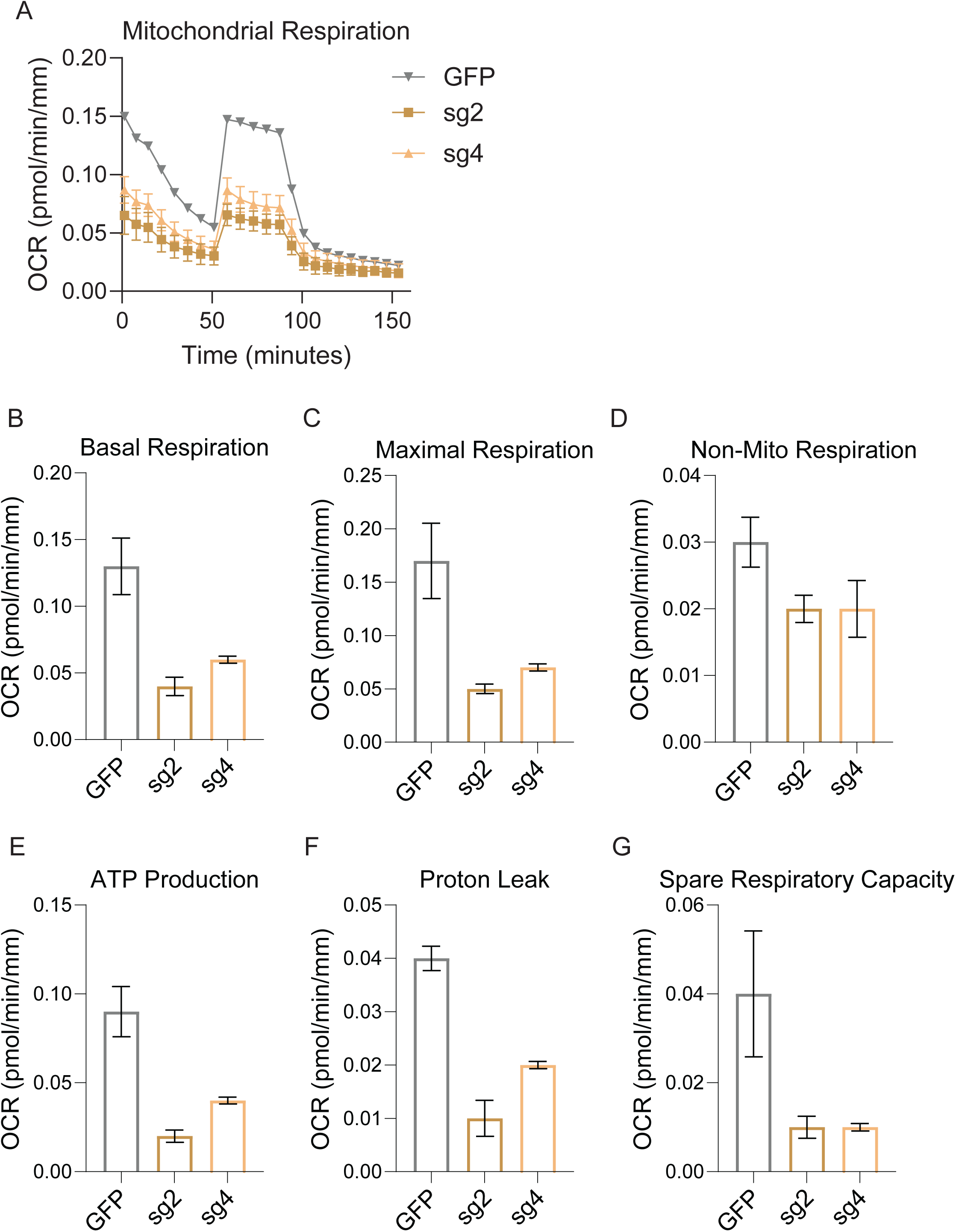
Mito Stress Test Assay **(A)** raw and **(B)** normalized OCR kinetic graph of mitochondria respiration in wildtype (WT) and knockout (sg2 and sg4) pediatric brain tumor neurospheres. OCR bar chart of each bioenergetic measurement, including **(C)** basal respiration, **(D)** maximal respiration, **(E)** non-mitochondria respiration, **(F)** ATP production, **(G)** proton leak, **(H)** spare respiratory capacity.

## Resource availability

Technical contact: Technical questions on executing this protocol should be directed to and will be answered by the technical contact, Jessica Tsai

This study did not generate new unique reagents.

## Acknowledgments

Research funding to J.W.T: National Cancer Institute 1K08CA279908-01A1, Rally Foundation, Kids Join the Fight, St. Baldrick’s Foundation, Griffin’s Guardians, Alex’s Lemonade Stand Foundation, the Wright Foundation, Musella Foundation for Brain Tumor Research, Stache Strong, the Albert and Bettie Sacchi Foundation, McKenna Claire Foundation, the Elsa U. Pardee Foundation, the Margaret E. Early Medical Research Trust, Wipe Out Kids’ Cancer, The Cure Starts Now Foundation, Brooke Healey Foundation, Melina Michelle Edenfield Foundation, The Cure Starts Now Australia, The Cure Starts Now Canada, Reflections Of Grace Foundation, Yuvaan Tiwari Foundation, Cure Brain Cancer Foundation, Aubreigh’s Army Foundation 328, Aidan’s Avengers, Run DIPG, Musella Foundation, Love4Lucas Foundation, Whitley’s Wishes, Anna’s Bake Sale Foundation, The Ayla Foundation, The Isabella and Marcus Foundation, Love, Chloe Foundation, Lauren’s Fight for Cure, Robert Connor Dawes Foundation, Ryan’s Hope, The Gold Hope Project, Abby’s Corner Foundation, the DIPG/DMG Collaborative, and Snapgrant.com. We are also thankful to the Cancer and Blood Disease Institute (CBDI) at Children’s Hospital Los Angeles (CHLA), the Saban Research Institute at CHLA, and USC Norris Comprehensive Cancer Center for institutional startup funds.

## Author contributions

S.T.: Conceptualization, data collection, data analysis, writing, editing. J.W.T.: Conceptualization, writing, editing, supervision, funding acquisition.

